# GLUE: A flexible software system for virus sequence data

**DOI:** 10.1101/269274

**Authors:** Joshua B Singer, Emma C Thomson, John McLauchlan, Joseph Hughes, Robert J Gifford

## Abstract

**Background:** Virus genome sequences, generated in ever-higher volumes, can provide new scientific insights and inform our responses to epidemics and outbreaks. To facilitate interpretation, such data must be organised and processed within scalable computing resources that encapsulate virology expertise. GLUE (Genes Linked by Underlying Evolution) is a data-centric bioinformatics environment for building such resources. The GLUE core data schema organises sequence data along evolutionary lines, capturing not only nucleotide data but associated items such as alignments, genotype definitions, genome annotations and motifs. Its flexible design emphasises applicability to different viruses and to diverse needs within research, clinical or public health contexts.

**Results:** HCV-GLUE is a case study GLUE resource for hepatitis C virus (HCV). It includes an interactive public web application providing sequence analysis in the form of a maximum-likelihood-based genotyping method, antiviral resistance detection and graphical sequence visualisation. HCV sequence data from GenBank is categorised and stored in a large-scale sequence alignment which is accessible via web-based queries. Whereas this web resource provides a range of basic functionality, the underlying GLUE project can also be downloaded and extended by bioinformaticians addressing more advanced questions.

**Conclusion:** GLUE can be used to rapidly develop virus sequence data resources with public health, research and clinical applications. This streamlined approach, with its focus on reuse, will help realise the full value of virus sequence data.

## Background

The study of virus genome sequences is important in medical, public health, veterinary and basic research contexts. Recent advances in sequencing technologies are driving a rapid expansion in the volume of available genomic sequence data for different viruses. Virus genome sequencing is now a key technology for understanding virus biology and for facing the challenges provided by viral outbreaks and epidemics.

To realise the full value of virus genome sequencing, sequence data must be processed within **virus sequence data resources**: scalable software systems that encapsulate the appropriate virology expertise. The components of these systems typically include both curated datasets and automated analysis, but these vary according to which species is targeted and the types of functionality offered. Table 1 shows some examples of well-established virus sequence data resources.

**Table 1.**
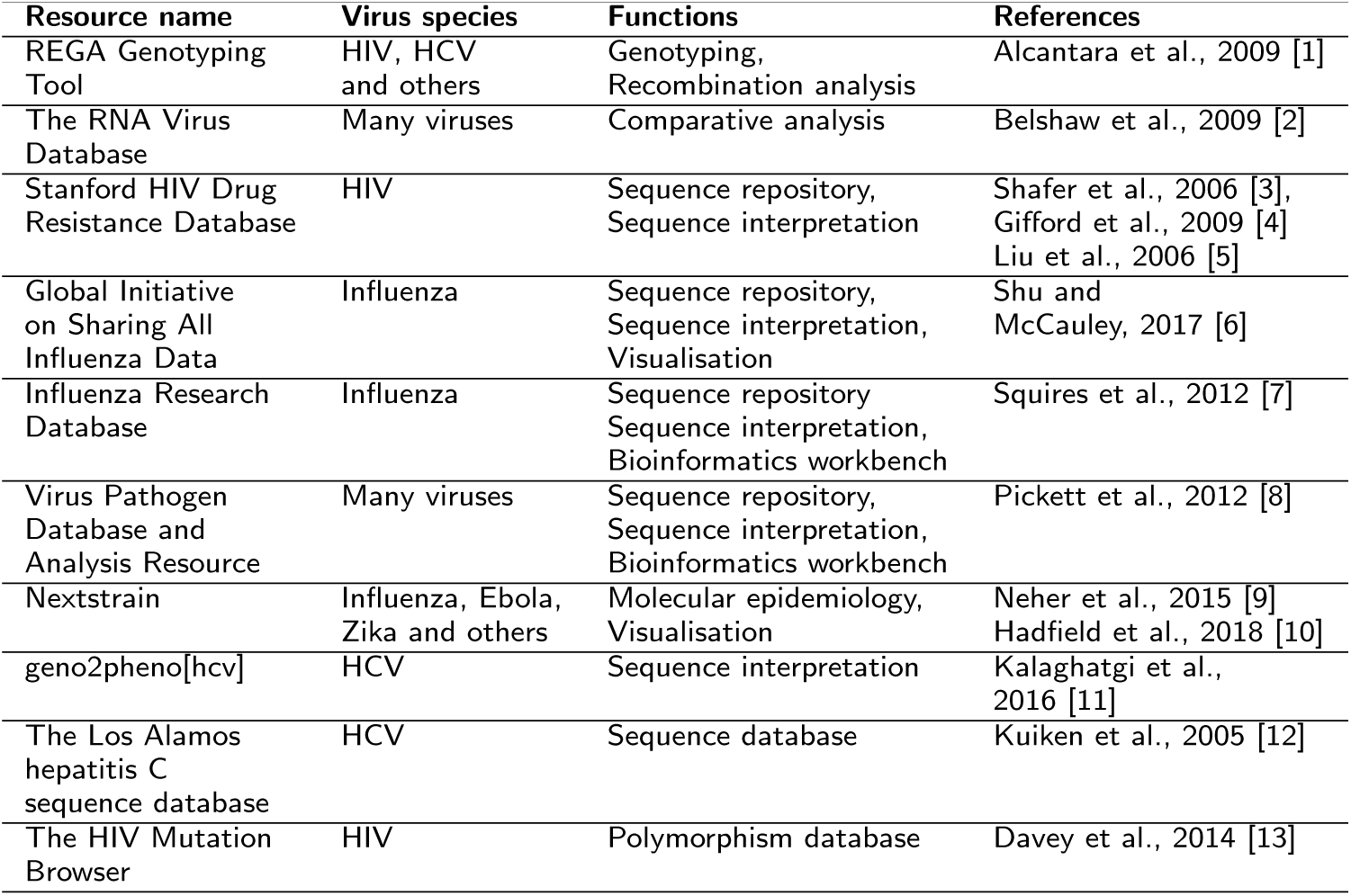
Examples of virus sequence data resources.

| Resource name | Virus species | Functions | References |
| --- | --- | --- | --- |
| REGA Genotyping Tool | HIV, HCV and others | Genotyping, Recombination analysis | Alcantara et al., 2009 [1] |
| The RNA Virus Database | Many viruses | Comparative analysis | Belshaw et al., 2009 [2] |
| Stanford HIV Drug Resistance Database | HIV | Sequence repository, Sequence interpretation | Shafer et al., 2006 [3], Gifford et al., 2009 [4], Liu et al., 2006 [5] |
| Global Initiative on Sharing All Influenza Data | Influenza | Sequence repository, Sequence interpretation, Visualisation | Shu and McCauley, 2017 [6] |
| Influenza Research Database | Influenza | Sequence repository, Sequence interpretation, Bioinformatics workbench | Squires et al., 2012 [7] |
| Virus Pathogen Database and Analysis Resource | Many viruses | Sequence repository, Sequence interpretation, Bioinformatics workbench | Pickett et al., 2012 [8] |
| Nextstrain | Influenza, Ebola, Zika and others | Molecular epidemiology, Visualisation | Neher et al., 2015 [9], Hadfield et al., 2018 [10] |
| geno2pheno[hcv] | HCV | Sequence interpretation | Kalaghatgi et al., 2016 [11] |
| The Los Alamos hepatitis C sequence database | HCV | Sequence database | Kuiken et al., 2005 [12] |
| The HIV Mutation Browser | HIV | Polymorphism database | Davey et al., 2014 [13] |

We have developed Genes Linked by Underlying Evolution (GLUE), a flexible software system for virus sequence data (http://tools.glue.cvr.ac.uk). The aim of GLUE is to facilitate the rapid development of diverse sequence data resources for different viruses. The GLUE “engine” is an open, integrated software toolkit that provides functionality for storage and interpretation of sequence data. The engine itself does not include any components specific to a particular virus. GLUE “projects”, on the other hand, capture data sets and other items relating to a specific group of viruses. These projects are hosted within the GLUE engine.

GLUE features several innovative aspects that differentiate it from existing work; (i) whereas most public virus sequence data resources do not make their internal software available, GLUE may be downloaded and used by anyone to create a new resource; (ii), GLUE is data-centric; all elements of a GLUE project are stored in a standard relational database, including sequence data, genome annotations, analysis tool configuration, and even custom program code; (iii) to manage high levels of variation within virus types, GLUE organises sequence data according to an evolutionary hypothesis, placing multiple sequence alignments at the centre of its design; (iv) the GLUE data schema is extensible, allowing projects to host auxiliary data, for example geographical sampling locations; (v) finally GLUE has a simple programmatic interface, which can be used not only in a conventional bioinformatics pipeline but also in web resources based on GLUE.

To test these ideas we developed HCV-GLUE, a sequence data resource for hepatitis C virus (HCV), briefly presented here as a case study. HCV-GLUE offers various data sets and computational functions including maximum likelihood phylogenetics and drug resistance analysis. Research bioinformaticians can download HCV-GLUE for use within their labs while an interactive web application (http://hcv.glue.cvr.ac.uk) provides sequence data, analysis and visualisation to a wider range of users.

GLUE provides a platform for the rapid development of powerful, reusable virus sequence data resources. These can inform virus research and also help us respond to challenges posed by existing and emerging viruses of concern.

## Implementation

### Scope

Virus sequence data resources vary along two major dimensions: the types of virus sequence which are of interest and the sequence data functionality which is offered.

A GLUE-based resource may relate to sequences from a single virus species, e.g. *human immunodeficiency virus 1*, or from some higher-ranked grouping, e.g. the family *Retroviridae*. It may encompass the whole genome of the viruses of interest or only a single segment or gene. In timescale terms it might cover only extant sequences from a single contemporary outbreak over a few months or years or, at the other extreme, hundreds of millions of years when the deeper evolutionary relationships between viruses are being examined, for example across lentiviruses infecting mammals [14].

A GLUE-based resource can fulfill a range of functions within research, public health, medical or veterinary contexts; any area where analysis of virus sequence variation is of value. One primary use of GLUE is to quickly build bespoke repositories for consensus nucleotide data. Sequence data may be derived from existing public datasets, as in the Influenza Research Database [7], or may be a product of research, clinical or public health activities. A key benefit of GLUE is that important genomic aspects of sequences, such as protein translations of specific genome regions, may be quickly computed and made widely accessible. In clinical scenarios, this allows improved analysis of viral infections, for example in detecting drug resistance of clinical relevance, as in HIVdb or geno2pheno[hcv] [5, 11].

Within GLUE, sequences can also be linked to any form of auxiliary data. Common examples of auxiliary data include the disease status of infected organisms as in the Los Alamos HCV database [12], and geographical or temporal data as in nextstrain [9, 10]. GLUE can therefore serve as a bioinformatics platform for investigating relationships between genomic variation and these other variables. In public health, a GLUE-based resource with epidemiology-related metadata may play a role in real-time molecular surveillance as suggested in [15].

Sequence datasets can be combined with analysis functionality in an integrated GLUE project. A core project for a virus species would typically define a set of reference sequences, basic genome features and the important agreed clades within the species. It may also provide clade assignment functionality, as in the REGA genotyping tools [1]. The project can then be disseminated for example using public version control repositories [16]. Publication of such a core project can then promote standards for organising and analysing sequence data and allows the community of interest to avoid recapitulating the basic tasks of sequence data organisation for the virus each time sequence analysis projects are undertaken.

Extensions to a core project can add more specialist data and new analysis modules. One direction for a project extension is to catalogue specific genomic variations such as amino acid polymorphisms, as in HIVmut [13], including drug resistance as in HIVdb or geno2pheno[hcv] [3, 11] or epitopes as in IEDB [17].

The value contained within GLUE projects can be leveraged in a wide variety of computing contexts. To facilitate this, GLUE relies on a minimal set of mature, high-quality, cross-platform software components. GLUE can import and export data in various formats and contains powerful scripting and command line capabilities which allow it to be quickly integrated into a wider bioinformatics environment. GLUE may also be deployed within a standard web server, allowing its functionality to be exposed via standard web service protocols for machine-to-machine interaction. This capability can be used to build interactive public websites or public programmatic services, or to integrate GLUE into the wider computing infrastructure of an organisation as part of a microservices software architecture.

### Design overview

The GLUE engine is the software package on which all GLUE-based virus sequence data resources depend. Its features are intended to be useful across a broad range of GLUE-based resources without being specific to any virus or usage scenario. Interaction with the GLUE engine is mediated via the **command layer**, its public interface. The command layer can be used to manipulate and access stored data, and to extend the data schema associated with a project. It also provides a range of fine-grained bioinformatics functions along with mechanisms for adding custom analysis functionality.

A GLUE project is essentially a dataset focused on a particular virus and/or analytical context and held within the GLUE data schema. Project development requires the collation and curation of heterogeneous data: sequences, metadata, genome annotations, clade definitions, alignments, phylogenetic trees and so on. Command scripts are then used to load project data into the GLUE database and integrate it using relational links. A project may be further extended with configuration of functionality such as clade assignment methods, the design of data schema extensions and the implementation of any custom analysis functionality.

This separation of concerns has a range of benefits. GLUE project developers focus on using the GLUE command layer to develop their projects. So, while GLUE projects depend on the syntax and semantics of the command layer they do not depend directly on the internal details of the GLUE engine, which is implemented in Java. System-wide aspects such as database access and schema management, interfacing with bioinformatics software and the provision of web services are handled by the GLUE engine. GLUE-based resources benefit from these without significant effort from project developers.

GLUE projects may be hosted in local repositories or cloud-based public or private repositories such as GitHub [16], allowing controlled collaborative development of these resources. Individual GLUE projects are version-controlled separately from the GLUE engine and from each other, so that individual projects can be maintained at a readily comprehensible scale.

### Data-centric architecture

GLUE has a data-centric, model-driven architecture. It defines a data schema and set of functions that support the common requirements of diverse virus sequence data resources. All information required for sequence processing, including both data and analysis configuration, is stored in a standard relational database structured according to this schema. Functions retrieve the required information from the database as required during any computation. There are several benefits to this approach.

Standard database mechanisms such as structured queries, relational joins, paging and caching may be employed, simplifying the implementation of higher-level logic. Cross-cutting concerns are simplified. For example referential integrity validation, query syntax and data exporting are all handled in a uniform way. Finally, the deployment of a GLUE-based resource on a new computer system is as simple as installing GLUE and then copying across the database contents; this is sufficient to ensure that all required data and analysis functionality is in place.

The GLUE **core schema** (Figure 1) is a fixed set of object types and relationships available in every GLUE project. The schema brings a certain level of evolution-oriented organisation to virus sequence data and captures the objects and relationships most commonly required to utilise them.

**Figure 1.**
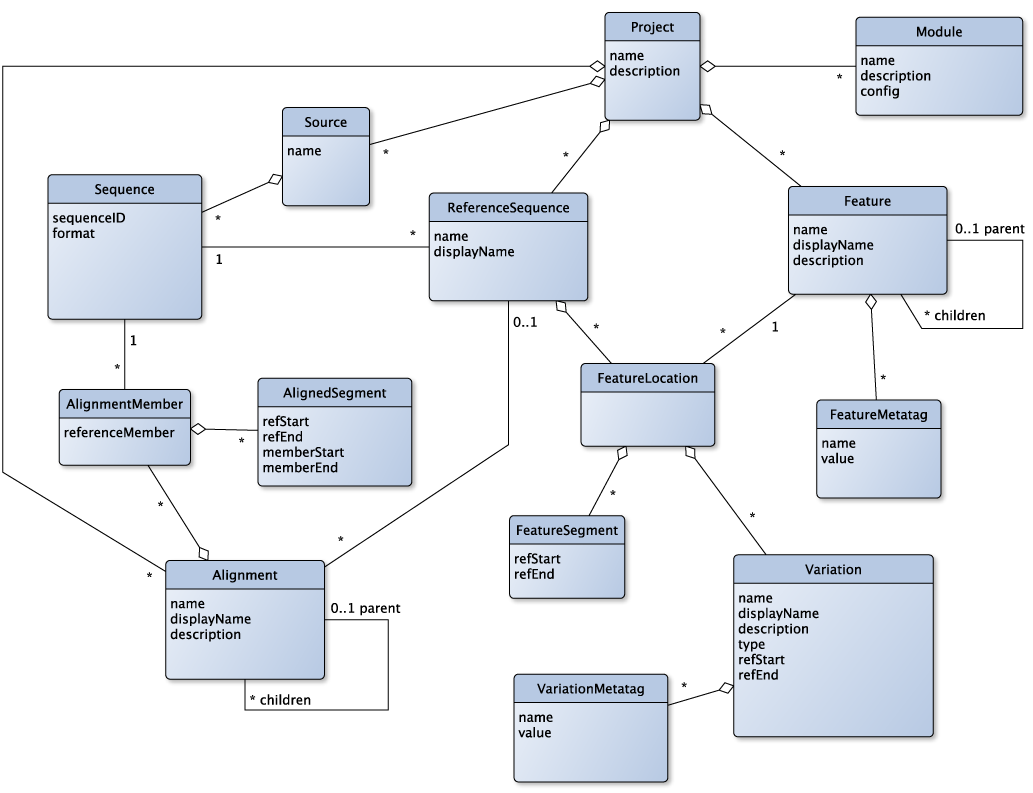
The GLUE core schema. The main object types, fields and relationships in the core schema of GLUE, represented as a Unified Modelling Language (UML) entity-relationship diagram. Object types are represented as blue boxes with fields specified inside. Relationships between types are represented as lines connecting the associated object types. A diamond indicates a composition relationship. Relationship lines are annotated with “multiplicities” indicating how many objects participate in a single instance of the relationship; an asterisk indicates any number of objects.

The design is model-driven in the sense that the semantics of the core schema reflect concepts inherent to virus biology. This knowledge capture approach is similar to systems such as Gene Ontology [18, 19], Sequence Ontology [20] and Chado [21]. However, the set of concepts in the GLUE core schema is small and targeted at the requirements of virus sequence data resources. Furthermore, the semantics are flexible and intended to be adapted on a per-project basis, in contrast with these more formal ontologies. In presenting the core schema, we will first discuss why sequence alignments are central in its design, and then outline the main object types and their semantics.

#### The role of sequence alignments

Nucleotide-level multiple sequence alignments are hypotheses about evolutionary homology and are critical for interpreting virus sequence data. The pairwise alignment of a new sequence with a well-understood existing sequence allows the location of genomic features within the new sequence to be inferred. The construction of multiple sequence alignments allows more complex comparative analyses to be performed. For example, comparative approaches can be used to investigate properties such as phylogenetic relationships, selection pressures, and evolutionary conservation.

The creation of high-quality alignments can require a significant investment of effort. Distinct virus genome regions or sets of sequences may require different techniques, for example alignment of distantly related sequences typically requires a degree of human oversight, whereas closely related sequences may be reliably aligned automatically. Nucleotide alignments for coding regions are best performed with the knowledge of the translated open reading frame. The process of creating alignments also has a complex interdependency with tree-based phylogenetics: an alignment is a prerequisite for running a phylogenetics method and yet the incorporation of a new sequence into an alignment is strongly informed by the phylogenetic classification of that sequence.

Because alignments are critical to virus sequence data resources, GLUE places these high cost and high value resources at the centre of its strategy for organising sequence data. A key aim of the GLUE core schema is to capture as much nucleotide homology as possible amongst the sequences of interest, and to integrate it into a single data structure.

GLUE object types are denoted in italicised CamelCase, e.g. *FeatureLocation*, and their fields are denoted by lower case italics with angle brackets, e.g. *<sequenceID>*.

#### Sequences and Sources

As discussed above, viral nucleotide sequences form the foundation of a GLUE project. Each GLUE *Sequence* object is a viral nucleic acid, RNA or DNA. *Sequences* may originate from a variety of methodological approaches as long as consensus nucleotide strings are produced. A set of *Sequences* is grouped together within a *Source* object: *Sequences* are identified by the *Source* to which they belong, together with a *<sequenceID>* field that is unique within the *Source*.

#### Features

*Feature* objects represent parts of the viral genome with established biological properties. Coding *Features* are introduced for regions that are translated into proteins; there may also be *Features* for non-coding promoters, untranslated regions, introns and others. *Features* may be arranged in a hierarchy, reflecting the containment relationships of the corresponding genome regions (e.g. specific domains within a protein, or individual proteins within a precursor polyprotein).

#### ReferenceSequences and FeatureLocations

GLUE uses *ReferenceSequence* objects to organise, link and interpret sequence data within a project. A *ReferenceSequence* is based on a specific *Sequence*. The choice of which *Sequence* objects to use for *ReferenceSequence* objects can vary based on conventions within the virus research field or pragmatic concerns. *ReferenceSequences* contain *FeatureLocation* objects that provide specific co-ordinates for *Features*. Typically, multiple *ReferenceSequences* will contain *FeatureLocations* for the same *Feature*, but with different co-ordinates as necessary. Additionally, a certain *Feature* may be represented by *FeatureLocations* on a subset of *ReferenceSequences* since a certain gene for example may be present in the genomes of only certain viruses within the project.

#### Alignments and AlignmentMembers

Evolutionary homology proposes that a certain block of nucleotides in one sequence has the same evolutionary origin as a certain block in another sequence. *Alignment* objects aggregate statements of evolutionary homology between *Sequences*. An *Alignment* contains a set of *AlignmentMember* objects; each *AlignmentMember* associates a member *Sequence* with the containing *Alignment*. Each *Alignment* has a reference coordinate space and the *AlignmentMembers* contain *AlignedSegment* objects representing statements of homology within this space. Each *AlignedSegment* has four integer fields: *<refStart>*, *<refEnd>*, *<memberStart>* and *<memberEnd>*. An *AlignedSegment* states that the nucleotide block *<memberStart>*:*<memberEnd>* in the member *Sequence* is to be placed at location *<refStart>*:*<refEnd>* in the reference coordinate space of the *containing Alignment*. This indirectly relates member sequence nucleotides with each other: blocks of nucleotides from distinct *Sequences* are homologous within an *Alignment* when they are placed at the same reference coordinate location.

*Alignment* objects in GLUE are data structures which store nucleotide homologies between sequences. The possible sources of these homologies include popular computational methods such as MAFFT [22] but also manual techniques.

There are two types of GLUE *Alignment*. In “unconstrained” *Alignments* the reference coordinate space is purely notional; not based on any particular *Sequence*. Nucleotide position columns in this coordinate space may be added in an unrestricted way in order to accommodate any homology between member *Sequences*.

A “constrained” *Alignment* is associated with a “constraining” *ReferenceSequence*. This provides a concrete coordinate space for the *Alignment*. *AlignedSegment* objects within a constrained *Alignment* propose a homology between a nucleotide block on a member *Sequence* and an equal-length block on the constraining *ReferenceSequence*.

Unconstrained *Alignments* have the advantage of being able to represent the full set of homologies between any pair of member sequences. However they must utilise an artificial coordinate space to achieve this, and this coordinate space must expand to accommodate every insertion, potentially leading to a large, unmanageable set of columns. Conversely, constrained *Alignments* use a fixed, concrete coordinate space but cannot represent homologies for nucleotide columns contained within insertions relative to the constraining *ReferenceSequence*.

#### Variations

Patterns of residues within virus sequences, at both the nucleotide and amino acid levels, are associated with specific functions or phenotypes. Knowledge about such residue patterns is typically derived from testing specific laboratory-derived or - modified virus strains in specific assays or observing their specific phenotypes. As these patterns become more established in the literature, it is informative to investigate the extent to which they may be present in a broader set of related viruses. A *Variation* is a named nucleotide or amino-acid residue pattern. *Variations* may also describe insertions or deletions at the nucleotide or amino-acid level. GLUE contains functions to analyse *Variations* in sequence data.

Patterns associated with a *Variation* may be defined as concrete strings of nucleotides or amino acids. Alternatively, for greater expressive flexibility, regular expressions may be used. These are patterns to be matched within a target string, with a standardised syntax and semantics. The biological properties of a *Variation* pattern may be captured via a data schema extension as discussed in the following section.

Each *Variation* is contained within a *FeatureLocation* object belonging to a specific *ReferenceSequence*, in order to anchor it to a genomic location. This allows documented residue patterns from the research literature to be quickly incorporated into a GLUE project using standardised reference coordinates.

#### Schema extensions

Virus sequence resources often contain important auxiliary data items which cannot be captured within the GLUE core schema. These project-specific data objects may have highly structured relationships with objects in the core schema and with each other. GLUE provides a powerful yet easy-to-use mechanism for extending the data schema on a per-project basis. New fields may be added to tables in the core schema. New custom object types may be added, with their own data fields. Finally, custom links may be added between pairs of object types in the schema.

For example *Rabies lyssavirus* is a negative-sense single-stranded RNA virus in the *Rhabdoviridae* family infecting a variety of animal species including humans. The wide host range of this virus suggests that a GLUE project for the virus may need to represent the host species from which each viral sequence was originally obtained. A custom object type can be introduced with an object for each possible host species. A custom many-to-one link can associate each viral sequence with the host species from which it was sampled. Host species objects themselves could then be annotated with ecological factors or associated with host genus or host family objects if these higher-rank taxonomic groups are of interest.

#### Object-relational mapping

Object-relational mapping (ORM) is a standard technique which allows application software to use object-oriented constructs such as classes, objects, fields and references to query and manipulate relational database entities such as tables, rows, columns and relationships. Internally, GLUE uses Apache Cayenne [23] as its ORM system. Data items from the core schema or extensions are represented as objects with fields and relationships, providing a convenient abstraction for GLUE commands and scripting logic. One example where this abstraction may be used is in filter logic supplied to GLUE commands. If the host species schema extension mentioned above is in place, the list sequence command may use a --whereClause filter option written in Cayenne’s expression language to request all sequences where the host species is, for example, within the family Canidae:

~~~
list sequence --whereClause "host species.family.id = ‘Canidae’"
~~~

This applies a filter to the *Sequence* table, requiring each selected object to be associated with a host species object, which is in turn associated with the host family object with ID “Canidae”. The filter logic is expressed in object-oriented terms, but translated into SQL JOIN syntax internally.

### The alignment tree

GLUE projects have the option of using a structure called an “alignment tree”, which links together nucleotide homologies in an evolution-oriented way. The alignment tree captures established evolutionary relationships between sets of sequences and integrates these with nucleotide-level homology data.

An alignment tree is built by first creating constrained *Alignment* objects for each of the established clades for the viruses of interest. Where a parent-child relationship between two clades exists within the evolutionary hypothesis, a special relational link is introduced between the corresponding pairs of *Alignment* objects. *Sequence* objects are then assigned to clades by adding them as *AlignmentMembers* of the corresponding *Alignment*.

It is important to note the processes by which clades within an evolutionary hypothesis are established. Homologies recovered from nucleotide sequence data offer a starting point for generating a detailed branching phylogenetic tree via a variety of computational methods, such as RAxML [24]. In using such methods, the intention is that this tree approximates the underlying evolutionary history. However, such techniques are subject to errors and uncertainties arising from various sources including the sequence sample set, the alignment and the choice of substitution model. Despite these limitations, some clear and robust phylogenetic evidence can emerge for clade relationships within a virus species as well as for clades at higher taxonomic ranks. The status of the evolutionary hypothesis concerning a set of viruses can therefore be at any point along a spectrum, depicted in stylised form in Figure 2. At one extreme, the “clade resolution” is minimal: the evolutionary history is unknown except that all sequences in the set belong to a single group. At intermediate points on the spectrum, some phylogenetic relationships between sequences remain unresolved, but virus sequences have been assigned to well-understood, widely-agreed clades, and the phylogenetic relationships between these clades have been established. At the far end of the spectrum, at the point of maximal clade resolution, a detailed phylogenetic tree has been established with each sequence on a leaf of the tree, and each internal node carrying a high degree of support.

**Figure 2.**
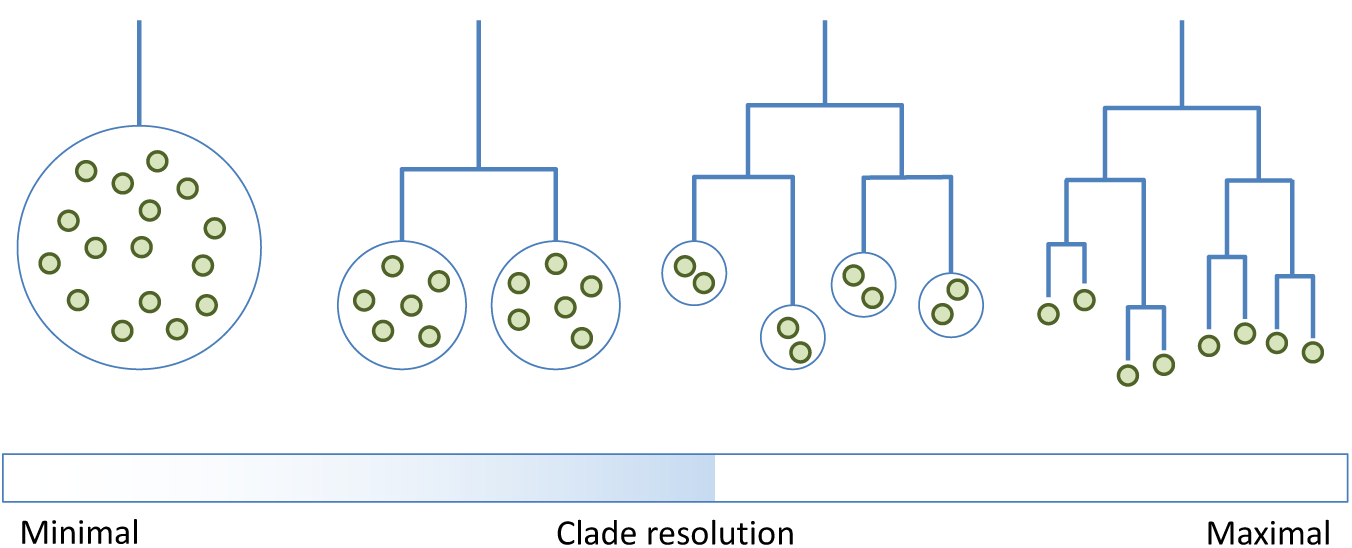
Different levels of clade resolution amongst viral evolutionary hypotheses. At minimal resolution, the set of viruses are known only to belong to the main clade. At maximal resolution, all phylogenetic relationships between viruses are known.

An alignment tree can represent the virus evolutionary hypothesis at any point along this spectrum. *Sequences* may remain as members of the same *Alignment* as long as their precise evolutionary relationship remains unclear. As the finer structure of the phylogeny emerges, perhaps as new strains are sequenced, new *Alignment* objects may be introduced to represent the newly-established clades, and sequences may be moved according to their clade assignment.

An invariant is a logical property of a software system which is always true. The GLUE engine enforces the “alignment tree invariant” in its operations on constrained *Alignments*: If *Alignment A* is a child of *Alignment B* the *Sequence* acting as the constraining *ReferenceSequence* of *Alignment A* must also be a member sequence of *Alignment B*. In this way, a parent *Alignment* is forced to contain representative member sequences from any child *Alignments*. The object structure of an example alignment tree, demonstrating the invariant, is shown in Figure 3.

**Figure 3.**
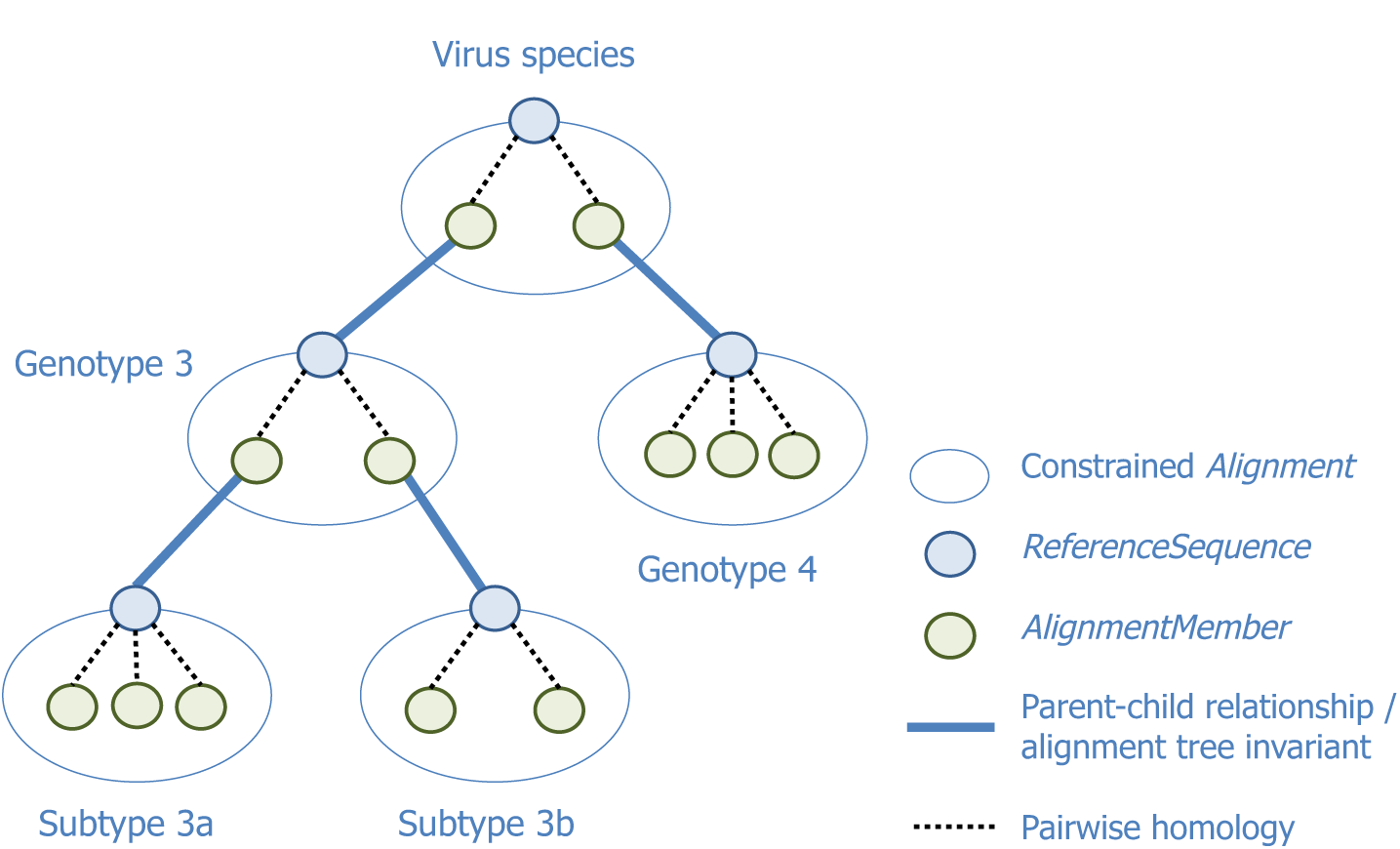
The object structure of an example alignment tree. The constrained *Alignment* at the root represents an entire virus species. Two child *Alignments* represent established clades: genotypes 3 and 4. Genotype 3 is further subdivided into two subtypes, 3a and 3b. Each constrained *Alignment* has a constraining *ReferenceSequence*. Within each *Alignment* node there are various *AlignmentMember* objects, each one records the pairwise homology between the member *Sequence* and the constraining *ReferenceSequence*. The alignment tree invariant requires for example that the constraining *ReferenceSequence* of subtype 3a is also a member of its parent, genotype 3.

In practice, *Alignments* at the tips of the tree contain the bulk of *Sequences*, as their memberships are determined by some clade assignment process. An *Alignment* at an internal position represents a putative ancestral clade, and only needs to directly relate together representatives of its descendent clades; sequences within these descendent clades are indirectly considered members of the ancestral clade. It may also be useful to place other sequences at an internal node if their membership of a more specific clade cannot be established.

As discussed, a constrained *Alignment* is unable to represent homologies which exist at positions within insertions relative to the *ReferenceSequence*. This is unlikely for member sequences of tip *Alignments* as long as the *ReferenceSequence* is a close relative. A group of sequences within an *Alignment* may contain an insertion relative to the *ReferenceSequence*. If the insertion contains data of interest to the project, this may warrant a new child *Alignment* with an appropriate constraining *ReferenceSequence*, containing the insertion. For *Alignments* at internal positions, one approach in future could be to use an ancestral reconstruction as the *ReferenceSequence*; consistent sets of insertions relative to this may then correspond to a new clade.

A significant advantage of the alignment tree is to fix the known evolutionary relationships between sequences and thereby avoid recomputing these in later analysis. The alignment tree also provides a pragmatic means to integrate different alignments computed using different techniques. Where sequences are closely related, reliable alignments can often be quickly built using simple pairwise methods to align sequences within an *Alignment* to the constraining *ReferenceSequence*, for example based on BLAST+ [25]. To obtain homologies for *Alignments* at internal positions where the relationship is more distant, manual or automated alignment methods, possibly operating at the protein level, may be more appropriate. In either case GLUE allows the corresponding nucleotide homologies to be imported and stored as *AlignedSegments* within the appropriate *AlignmentMember*.

Over the whole genome, two distantly-related virus sequences may be so divergent that it is impossible to fully align them reliably and analyse them together. However they may be much more conserved at specific genome regions. Internal or root nodes of the alignment tree can capture the homology for these conserved regions across a broad range of clades. Alignment tree nodes nearer to the tips may capture homology for a larger fraction of the genome, but for a narrow range of clades.

The alignment tree invariant guarantees that between any two *Sequence* objects, there is a path of *AlignmentMember* associations and corresponding pairwise sequence homologies. A simple transitivity idea composes homologies along the path into a single homology. For example if nucleotide block 21:50 on sequence *A* is homologous to block 31:60 on sequence *B*, and block 41:70 on sequence *B* is homologous to block 1:30 on sequence *C*, then block 31:50 on *A* is homologous to block 1:20 on *C*. GLUE applies this technique in various situations which require a homology between *Sequences* in different *Alignments* within the tree.

### The command layer

The command layer forms the programmatic interface of the GLUE engine. Commands cover a range of fine-grained functions including the manipulation and querying of any element in the project data set or the project schema extensions. Other commands perform more high-level functions; some examples are listed in Table 2.

**Table 2.**
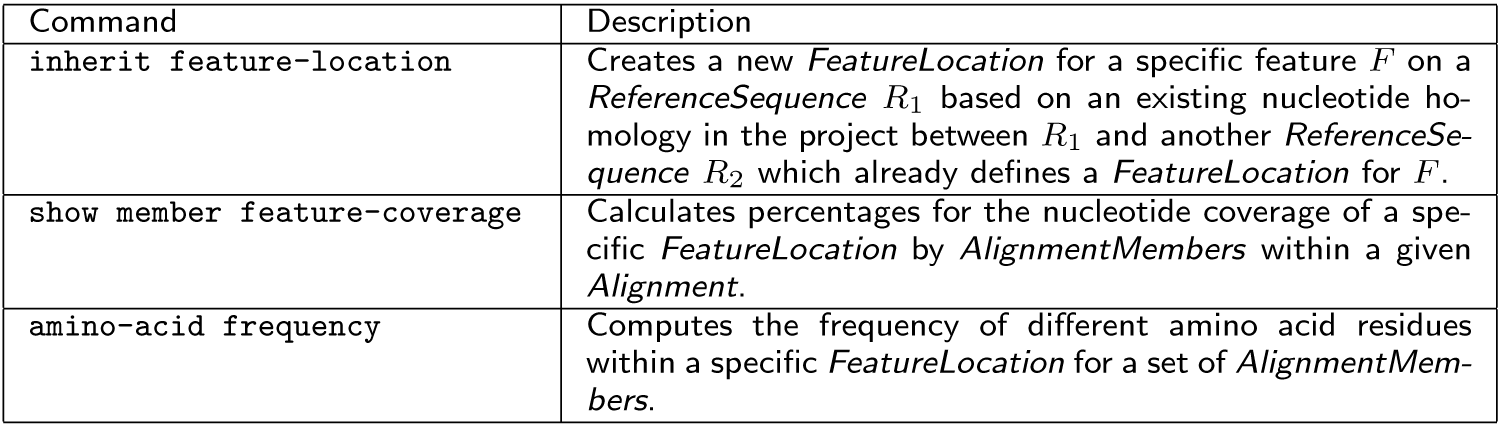
Examples of GLUE commands with high-level functions.

| Command | Description |
| --- | --- |
| inherit feature-location | Creates a new *FeatureLocation* for a specific feature *F* on a *ReferenceSequence* *R*<sub>1</sub> based on an existing nucleotide homology in the project between *R*<sub>1</sub> and another *ReferenceSequence* *R*<sub>2</sub> which already defines a *FeatureLocation* for *F*. |
| show member feature-coverage | Calculates percentages for the nucleotide coverage of a specific *FeatureLocation* by *AlignmentMembers* within a given *Alignment*. |
| amino-acid frequency | Computes the frequency of different amino acid residues within a specific *FeatureLocation* for a set of *AlignmentMembers*. |

A significant amount of functionality in the command layer is provided via the “module” mechanism. The current release of the GLUE engine provides more than 40 module types (documented online). When a module is created, commands associated with the module type become available. Modules are stored data objects, each module contains a configuration document which modulates the operation of the module commands, for example providing a set of rules or numeric parameter settings. In this way built-in functionality can be adapted on a per-project basis. GLUE module commands perform a wide variety of functions and can include any use of or update to the project dataset, operations on data obtained from the local file system or attached to an incoming web request, and operations involving external bioinformatics programs such as BLAST+. Examples of module types are given in Table 3.

**Table 3.** Examples of GLUE module types.

| Module type | Description |
| --- | --- |
| `fastaProteinAlignmentExporter` | Creates amino-acid level alignments from *Alignments* within the GLUE project. A protein-coding *FeatureLocation* is specified along with a set of *AlignmentMember* objects selected from a given *Alignment* and its descendants. A translated amino-acid alignment is generated based on the stored homologies for the selected member sequences, which can then be exported to a file or used in further computation. |
| `blastProteinFastaAlignmentImporter` | Imports amino-acid level alignments into the GLUE project to be stored as nucleotide alignments. For each row of this input alignment a GLUE *Sequence* object is identified. TBLASTN is used to compare the alignment row with the nucleotides of the identified sequence. In this way, the nucleotide-level homologies implicit in the file are identified and *AlignedSegment* objects representing this homology are created within an unconstrained GLUE *Alignment*. |
| `ncbiImporter` | Runs an eSearch query on the GenBank database [26, 27], based on a configurable search term. Records are downloaded in GenBank XML format and stored as GLUE *Sequence* objects. |
| `genbankXmlPopulator` | Operates on a set of *Sequence* objects which are stored in GenBank XML format. According to configurable rules, it extracts data items from the GenBank XML, executes transformations on them and updates auxiliary data fields or associations on the corresponding *Sequence* object. |
| `samReporter` | Provides functionality for interpreting SAM/BAM files [28] containing high throughput sequencing data. One example is the amino-acid command, which will translate those reads in the file which map to a specific protein-coding feature in the project. The command outputs the proportions of amino acid residues found at each location. |

#### The command layer in use

GLUE contains an interactive command line environment focused on the development and use of GLUE projects by bioinformaticians. This provides a range of productivity-oriented features such as automatic command completion, command history and interactive paging through tabular data. It could be compared to interactive R or Python interpreters [29, 30], or command line interfaces provided by relational database systems such as MySQL [31].

A GLUE-based resource may have project-specific analysis or data manipulation requirements. For example the assembly of the alignment tree set may need to iterate over a certain set of clades to process each associated tip alignment. Analysis logic may need to extract alignment rows from the data set and compute a specific custom metric for each row. To address such requirements GLUE project developers may write JavaScript programs, based on the ECMAScript 5.1 standard [32]. These programs may invoke any GLUE command, and access the command results as simple JavaScript objects. The programs may then be encapsulated as GLUE modules with their code stored in the project database. They provide functionality to higher level code in the form of module commands.

Web services have become a *de facto* standard for machine-to-machine interaction, using HyperText Transfer Protocol (HTTP) to carry JavaScript Object Notation (JSON) or eXtensible Markup Language (XML) requests and responses between computer systems. A software resource may offer its application programming interface (API) as a web service to allow integration with other systems either over the public web or on a private network as part of a microservices architecture. GLUE may be embedded in a standard web server. In this case its command layer becomes accessible as a web service. Commands are sent as JSON documents attached to an HTTP POST request, using a uniform resource locator (URL) identifying a data object within a GLUE project. The command result document is encoded as JSON attached to the HTTP response.

### Maximum-likelihood clade assignment

The assignment of a set of sequences to the same clade asserts that they have a common ancestor which is more recent than any ancestor shared with a sequence assigned to an external clade. Maximum likelihood is a popular evaluation criterion for selecting an evolutionary tree to explain the origins of extant sequence data. As such it has played a strong role in studies which aim to identify clades with strong support [33]. Sequences may be assigned to clades using a simple similarity criterion, as in geno2pheno[hcv] [11]. While identity-based measures between sequences clearly do correlate with membership of real clades, maximum likelihood techniques provide a more principled methodology for placing sequences within an evolutionary hypothesis.

Building on existing maximum likelihood software, we have developed a new algorithmic method called Maximum Likelihood Clade Assignment (MLCA). An implementation of this MLCA is integrated into the GLUE engine. RAxML [24] is a highly optimised implementation of maximum likelihood phylogenetics. The core use of RAxML is to generate a full phylogenetic tree from a multiple sequence alignment. RAxML also contains a feature called the Evolutionary Placement Algorithm (EPA), which suggests high-likelihood branch placements for a new sequence on a fixed reference tree. EPA allows us to apply maximum likelihood without reconsidering the whole tree. In this sense EPA is well-suited to the problem of virus sequence clade assignment and forms the core of the MLCA method.

#### MLCA Overview

The role of MLCA is to assign one or more query sequences to clades defined in a reference dataset. Although in some contexts MLCA may be applied to batches of query sequences, it is important to note that MLCA computes a clade assignment for each query sequence individually, and does not perform any phylogenetic analysis aimed at relating query sequences within a batch to each other.

Clades can be defined at various phylogenetic levels. For this reason, we introduce the concept of a clade category. A clade category encapsulates a set of named clades which are mutually exclusive. Within a virus species, an example clade category would be “Genotype” which contains the major genotypes of the virus; Genotype 1, Genotype 2 etc.

#### MLCA Inputs

- A set of reference sequences *R*_1_*, R*_2_*, …*
- A multiple sequence alignment of these reference sequences
- A strictly bifurcating tree *T*_full_ with the reference sequences labelled at the tips of the tree.
- A set of named clade categories *C*_1_*, C*_2_*, …* and for each clade category, a set of clades. Each clade category additionally defines certain numeric parameters:
  – A distance cut-off *d*
  – A distance scaling exponent *s*
  – The final clade cut-off *f*
- For each clade *c*, a subtree *T*_c_ (i.e. internal node) of *T*_full_ is specified, which corresponds to this clade. The subtrees associated with the clades within a clade category must be mutually exclusive. The reference sequences which are leaf nodes of *T*_c_ are implicitly assigned to clade *c* (see figure 4).
- One or more query sequences *Q*_1_, *Q*_2_, …

**Figure 4.**
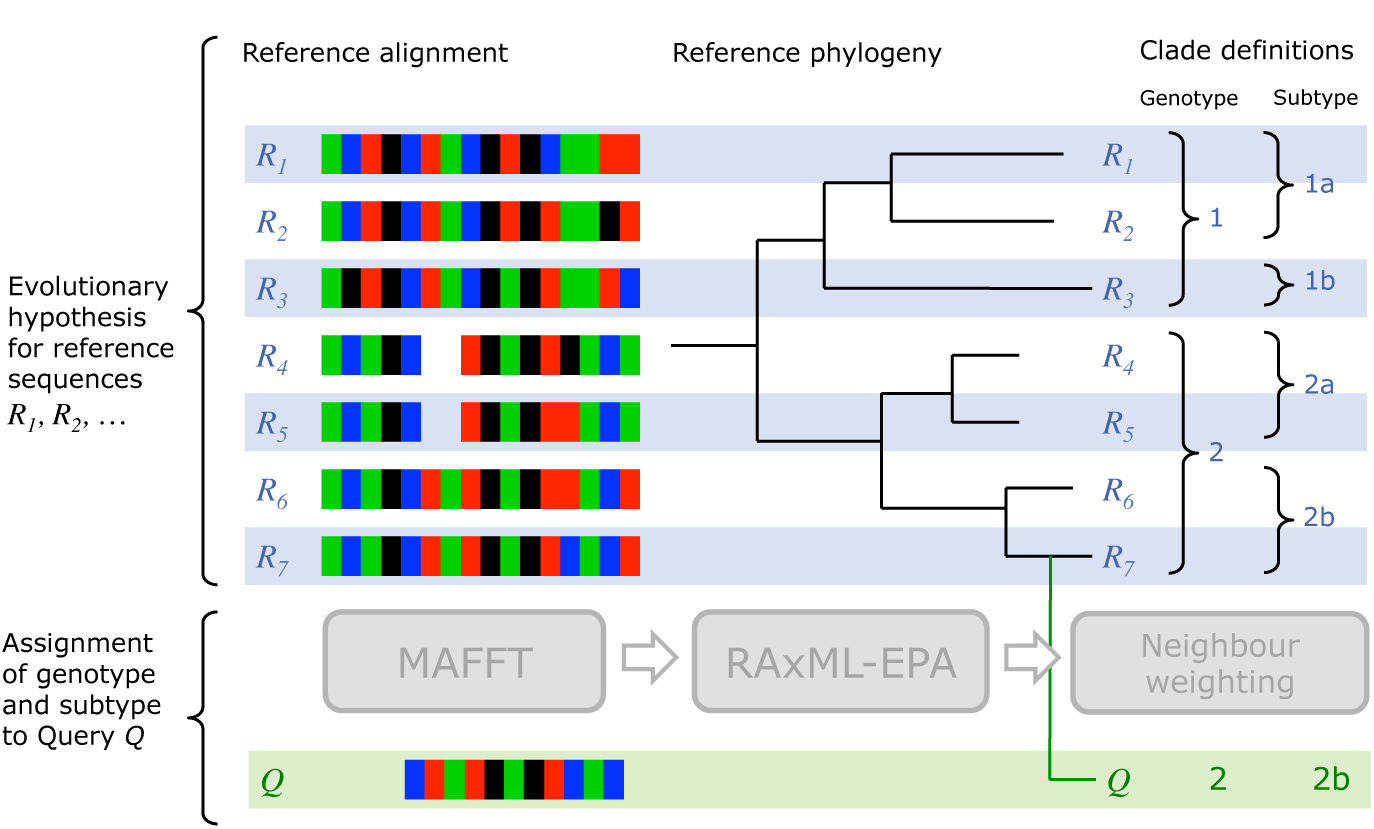
Graphical illustration of the MLCA algorithm. The evolutionary hypothesis for a virus within a GLUE project consists of an alignment of reference sequences *R*_1_, …, *R*_7_, a reference phylogeny with these sequences as leaf nodes and a set of clade definitions. In its initial alignment step, MLCA extends the reference alignment with a row for query sequence *Q*. Next, the placement step (RAxML EPA) suggests a branch for *Q* within the reference phylogeny. Finally, the Neighbour-weighting step assigns clades to *Q* by analysing the location of the additional branch in relation to neighbouring reference sequence taxa.

#### MLCA Outputs

- For each query sequence *Q*, a (possibly empty) set of strictly bifurcating trees, each tree consisting of *T*_full_ plus one additional branch for query *Q*, placed anywhere within *T*_full_.
- For each query sequence *Q* and clade category *C*:
  – An assignment of sequence *Q* to one of the clades in category *C*, or possibly no such assignment.
  – Percentage weights assigned to a subset of the member clades of *C*, or possibly an empty set of percentage weights.

#### MLCA Algorithm

The MLCA algorithm has three stages: alignment, placement and neighbour-weighting. A graphical illustration of the algorithm is presented in Figure 4.

The alignment stage takes as input the multiple sequence nucleotide alignment of the reference sequences *R*_1_*, R*_2_*, …* and produces as output an extended alignment which includes additional rows for the query sequences. In the GLUE implementation of MLCA, this step is based on the MAFFT software package [22]. We use the `--add` MAFFT option, which maintains the state of the initial alignment unchanged in the output and the `--keeplength` MAFFT option to avoid introducing additional nucleotide columns. A query sequence may include an insertion relative to the reference sequences but these are of no relevance to MLCA. Finally, if a batch of query sequences has been submitted to MLCA, we use a separate run of MAFFT for each query sequence, in order to keep the alignment computations for each query sequence independent.

The placement stage takes as input the extended alignment from the previous stage together with the tree *T*_*full*_. The aim of the placement stage is to find, for each query sequence *Q*, a set of placements *P*_1_*, P*_2_*, …*. Each placement *P* specifies a tree *T*_*P*_ comprising *T*_*full*_, plus a single additional branch for *Q*. The set of placements is selected such that the positions and lengths of the new branches maximise the likelihood of the extended tree given the extended alignment. The placement stage is implemented using the EPA subsystem of RAxML [24]. EPA operates by visiting each edge in *T*_*full*_ and inserting each query sequence at that edge in turn. After a query sequence is inserted, the insertion point and length of the new branch are optimised using RAxML’s maximum-likelihood infrastructure to find the maximum likelihood score possible for insertion at that edge. At the end of the process, a small set (possibly empty) of *n* high-likelihood placements *P*_1_*, P*_2_*, … P_n_* are retained for each query sequence.

Finally, the neighbour-weighting stage is summarises the outputs of the placement phase in the form of clade weightings and assignments. For a given query sequence *Q* and clade category *C*, neighbour-weighting will produce relative percentage weights for each clade *c* in *C*. Such a weight denoted *W_c,Q_* represents the strength of evidence that *Q* is a member of clade *c*. If the weight for a particular clade within *C* exceeds the final clade cut-off *f*, neighbour-weighting then recommends assigning *Q* to that clade.

The phylogenetic trees which RAxML produces use mean substitutions per site for branch lengths. The sum of branch lengths along the path between any two leaf nodes in the tree is the evolutionary distance between a pair of sequences, a measure of genetic relatedness.

This neighbour-weighting algorithm is based on the observation that a placement of a given query sequence *Q* implicitly ranks neighbouring reference sequences by increasing evolutionary distance from *Q*. Since each reference sequence *R* which neighbours *Q* is itself already assigned to a clade *c*, if *R* is a close neighbour, this contributes evidence to the assignment of sequence *Q* to clade *c*.

The pseudocode procedure CladeWeights in figure 5 specifies how the evolutionary distances to nearby reference sequences are combined to produce clade weights for a given query placement. The idea is modulated by applying *d* as a cut-off distance above which reference sequences are not considered to be a relevant neighbour. The *s* parameter is applied as a negative exponent to the evolutionary distance, allowing us to amplify the weight of nearer reference sequences.

**Figure 5.**
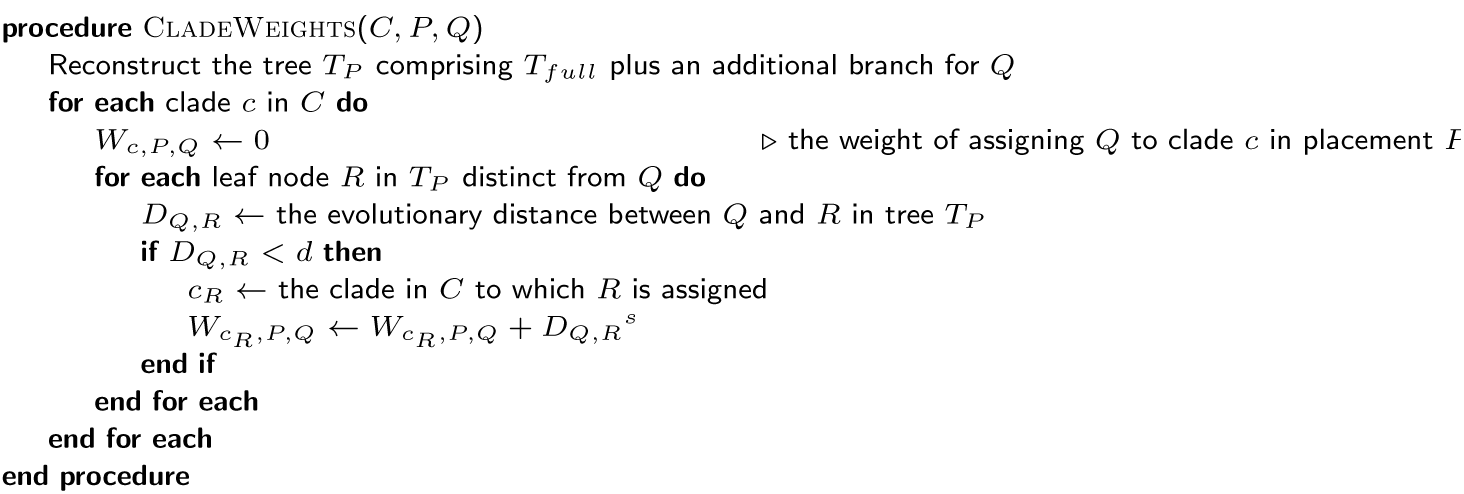
Procedure to assign clade weights for a query sequence. Given a placement *P* for a query sequence *Q*, a weight *W*_*c,P,Q*_ is calculated for each for clade *c* within a clade category *C*.

The EPA procedure used in the placement stage actually generates *n* placements *P*_1_*, P*_2_*, …, P_n_* for each query in the general case and also likelihood weight ratios *L*_1_*, L*_2_*, …, L_n_* for the placements such that these sum to 1 across the placement set. The neighbour-weighting stage combines the weights produced by CladeWeights across all placements within the set, to give a single weight for the query *Q* within each clade *c*. 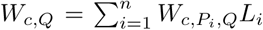. These weights are then normalised to percentages that sum to 100% across *C*. Finally *f* is used as a cut-off to determine whether a percentage weight is sufficient to recommend a clade assignment.

## Results

HCV-GLUE (http://hcv.glue.cvr.ac.uk) is a virus sequence data resource for hepatitis C virus (HCV) with both clinical and research applications. It is discussed here as a case study, to illustrate how GLUE-based resources may be developed. A rich set of web-based sequence data resources exists already for HCV [1, 8, 11, 12, 26]. HCV-GLUE complements these with a range of novel aspects.

Key items in the HCV-GLUE data set are taken from an accepted classification of HCV [33]. These items are the genotype and subtype clade definitions, a set of reference sequence definitions and an unconstrained master alignment of these reference sequences. From these items, an alignment tree is derived, with appropriate *ReferenceSequences* for each of the constrained Alignment nodes.

A large set of public HCV sequences is stored with associated metadata. These sequences are assigned to clades using a maximum-likelihood phylogenetic method, which is also made available in an online genotyping tool. The stored sequences are linked to each other via pre-built sequence alignments within the alignment tree.

The 5’ and 3’ untranslated regions are defined as GLUE *Features*, as are the open reading frame encoding the precursor polyprotein, which in turn has child *Features* for the 10 mature viral proteins derived from the polyprotein. GLUE provides commands for the analysis of amino acid residues within sequences; within HCV-GLUE these commands use the standardised numbering scheme proposed by Kuiken et al. [34].

HCV-GLUE provides online analysis of drug resistance for the direct-acting antiviral drugs (DAAs) that are available to treat HCV-infected patients. HCV-GLUE is the first sequence-based resistance analysis resource derived from European Association for the Study of the Liver (EASL) guidelines [35].

Besides the HCV-GLUE interactive website, the full dataset and analysis functions are available for any user to download and use on a private computer. The HCV-GLUE resource has been used in research articles: to characterise the public sequence data for drug target genes [36] and to predict the effectiveness of broadly neutralising antibodies which may play a role in a vaccine strategy [37].

### Clade Assignment

The MLCA method is implemented within HCV-GLUE and is used to assign genotypes and subtypes to sequences. The details of MLCA are described in the Implementation section.

Within HCV-GLUE, MLCA uses the polyprotein-coding region of the master unconstrained alignment. The reference phylogeny was generated from this alignment via a GLUE module that uses RAxML [24] with GTRGAMMA+I as the substitution model and 1000 bootstrap replicates. The placement step runs RAxML EPA with the GTRGAMMA+I substitution model. For the neighbour-weighting step Genotype and Subtype are defined as the two clade categories shown in Table 4.

**Table 4.** HCV-GLUE clade categories for MLCA. The Subtype category comprises 84 confirmed, 13 provisional and 42 unassigned subtypes, Also shown are category-specific numeric parameters for the neighbour-weighting step.

| Clade category | Number of clades | Distance cut-off $d$ | Distance scaling exponent $s$ | Final clade cut-off $f$ |
| --- | --- | --- | --- | --- |
| Genotype | 7 | 0.4 | -3.0 | 80% |
| Subtype | 139 | 0.26 | -3.0 | 80% |

We evaluated the MLCA method for HCV-GLUE by comparing its outputs against two other sources of clade assignments. One source of assignments was the GenBank metadata (referred to as Metadata); in many cases genotype and subtype have been supplied by the submitter of the sequence. A second source was the HCV genotyping procedure of the Virus Pathogen Database and Analysis Resource (ViPR [8]). This combines a distance-based phylogenetic tree computation approach with the branching index metric [38, 39] to produce genotype and subtype assignments.

The comparison was performed on the set of all GenBank sequences classified as HCV, of 500 bases or longer, downloaded on 19th December 2016. For each sequence we automatically extracted any genotype and subtype assignments from the Meta-data. We also obtained genotype and subtype results generated by the ViPR HCV genotyping method for each sequence. After patent-related sequences and known recombinants were excluded, there were 82,928 sequences in the comparison set.

Tables 5 and 6 show comparisons of assignments for the sequence genotype and subtype respectively. Each source of clade assignment may return a null result for genotype or subtype, and so the tables are separated into sections defined by the set of methods that returned non-null results. Within each section we report the number of sequences for which the different possible combinations of methods are in agreement.

**Table 5.** Comparison of HCV-GLUE genotype assignment with GenBank Metadata and assignment from ViPR.

| Methods giving non-null assignment results | Method agreement | Number of sequences |
| --- | --- | --- |
| Metadata & HCV-GLUE & ViPR | Metadata = HCV-GLUE = ViPR | 60 464 |
| | Metadata = HCV-GLUE ≠ ViPR | 518 |
| | Metadata ≠ HCV-GLUE = ViPR | 71 |
| | Metadata = ViPR ≠ HCV-GLUE | 0 |
| | Metadata ≠ HCV-GLUE ≠ ViPR | 2 |
|  | Total | 61 055 |
| Metadata & HCV-GLUE | Metadata = HCV-GLUE | 5019 |
| | Metadata ≠ HCV-GLUE | 26 |
|  | Total | 5045 |
| Metadata & ViPR | Metadata = ViPR | 6 |
| | Metadata ≠ ViPR | 7 |
|  | Total | 13 |
| HCV-GLUE & ViPR | HCV-GLUE = ViPR | 13 977 |
| | HCV-GLUE ≠ ViPR | 986 |
|  | Total | 14 963 |
| Metadata only | Total | 25 |
| HCV-GLUE only | Total | 1792 |
| ViPR only | Total | 4 |
| All assignment results null | Total | 31 |
|  | Grand total | 82 928 |

**Table 6.** Comparison of HCV-GLUE subtype assignment with GenBank Metadata and assignment from ViPR. A non-null subtype assignment implies a non-null genotype assignment.

| Methods giving non-null assignment results | Method agreement | Number of sequences |
| --- | --- | --- |
| Metadata & HCV-GLUE & ViPR | Metadata = HCV-GLUE = ViPR | 55 495 |
| | Metadata = HCV-GLUE ≠ ViPR | 1389 |
| | Metadata ≠ HCV-GLUE = ViPR | 1079 |
| | Metadata = ViPR ≠ HCV-GLUE | 47 |
| | Metadata ≠ HCV-GLUE ≠ ViPR | 4 |
|  | Total | 58 014 |
| Metadata & HCV-GLUE | Metadata = HCV-GLUE | 4508 |
| | Metadata ≠ HCV-GLUE | 206 |
|  | Total | 4714 |
| Metadata & ViPR | Metadata = ViPR | 7 |
| | Metadata ≠ ViPR | 13 |
|  | Total | 20 |
| HCV-GLUE & ViPR | HCV-GLUE = ViPR | 16 677 |
| | HCV-GLUE ≠ ViPR | 1306 |
|  | Total | 17 983 |
| Metadata only | Total | 129 |
| HCV-GLUE only | Total | 1667 |
| ViPR only | Total | 18 |
| All assignment results null | Total | 383 |
|  | Grand total | 82 928 |

For the majority of sequences, clade assignments were in agreement across all three methods; the proportion of such sequences was 72.91% for genotype and 66.92% for subtype. HCV-GLUE most frequently assigned a genotype (99.91%), followed by ViPR (91.69%) and finally Metadata (79.75%). Similarly, HCV-GLUE most frequently assigned a subtype (99.34%), followed by ViPR (91.69%) and finally Metadata (75.82%).

The higher frequency of a subtype assignment is partially explained by the fact that HCV-GLUE includes unassigned subtypes in its reference phylogeny and clade definitions. These are subtypes containing fewer than 3 distinct full-genome sequences and which have therefore not been given an official name. HCV-GLUE may assign a sequence to such a subtype. As of January 2018, there were about 200 such assignments, but HCV-GLUE has not yet revealed any unassigned subtype with 3 full-genome sequences.

In cases where all three methods assigned a genotype, there were two sequences where all methods disagreed. These were KJ678196 and KJ678224, from the same submission, both 822 bases in length. Aside from these, HCV-GLUE either agreed with the Metadata assignment or with ViPR.

For sequences where all methods assigned a subtype, there were 2 sequences where the methods agreed on the genotype but all disagreed on the subtype, these were AM401743, 576 bases in length and KP347290, 684 bases in length. Aside from these, HCV-GLUE agreed either with the Metadata assignment or with ViPR, with the exception of a group of 47 sequences. These sequences were derived from the same study [40], and each sequence was 543 bases in length. The sequence Metadata and ViPR assigned the sequences to subtype 1b whereas HCV-GLUE assigned them to subtype 1d. The HCV-GLUE assignment is plausible, for example geno2pheno[hcv] [11], which uses a BLAST-based approach, also assigns all 47 sequences to subtype 1d. Subtype assignments for shorter sequences such as these can be difficult to resolve.

The high level of agreement, at both genotype and subtype level, of HCV-GLUE assignments with ViPR and/or Metadata assignments supports the credibility of the HCV-GLUE clade assignment method.

### Public sequence data repository

HCV-GLUE acts as a downstream repository for HCV sequence data from GenBank, one of the principal public repositories for sharing virus sequence data [26, 27]. HCV-GLUE synchronises itself with GenBank HCV data on a daily basis, downloading any viral sequences of 500 bases or longer. HCV-GLUE adds value to the GenBank sequence set in several ways. Metadata is extracted from the GenBank XML format, normalised and added to custom columns of the HCV-GLUE *Sequence* table. Each sequence is then assigned a genotype and subtype where possible using the HCV-GLUE MLCA method and the sequence is added to the appropriate alignment tree node. A combined nucleotide and codon-aware BLAST-based method is used to generate a pairwise homology between the new sequence and the clade reference sequence.

The web application relies on the HCV-GLUE alignment tree as a navigational structure for accessing genotyped sequence data. Users may navigate to a specific clade and then apply simple web dialogs to filter sequences within the clade based on criteria including coverage of virus genome region, global region of origin and collection year.

Various data sets may then be downloaded based on the selected sequence set: a simple FASTA file of nucleotide data, metadata for the sequences in tabular format or a constrained multiple sequence alignment. The alignment may be composed of nucleotide or amino acid sequences and can be restricted to specific genome regions, for example mature proteins such as NS5A and sub-regions of these proteins.

### Automated genotyping and interpretation

The HCV-GLUE web application enables users to submit HCV nucleotide FASTA files for clade assignment and detailed sequence interpretation. Genotype and subtype assignments are presented to the user along with the closest neighbouring reference sequence and relative clade weightings produced by the MLCA procedure. Any atypical nucleotide or amino-acid polymorphisms are presented, along with a visual presentation of insertions and deletions (see Figure 6).

**Figure 6.**
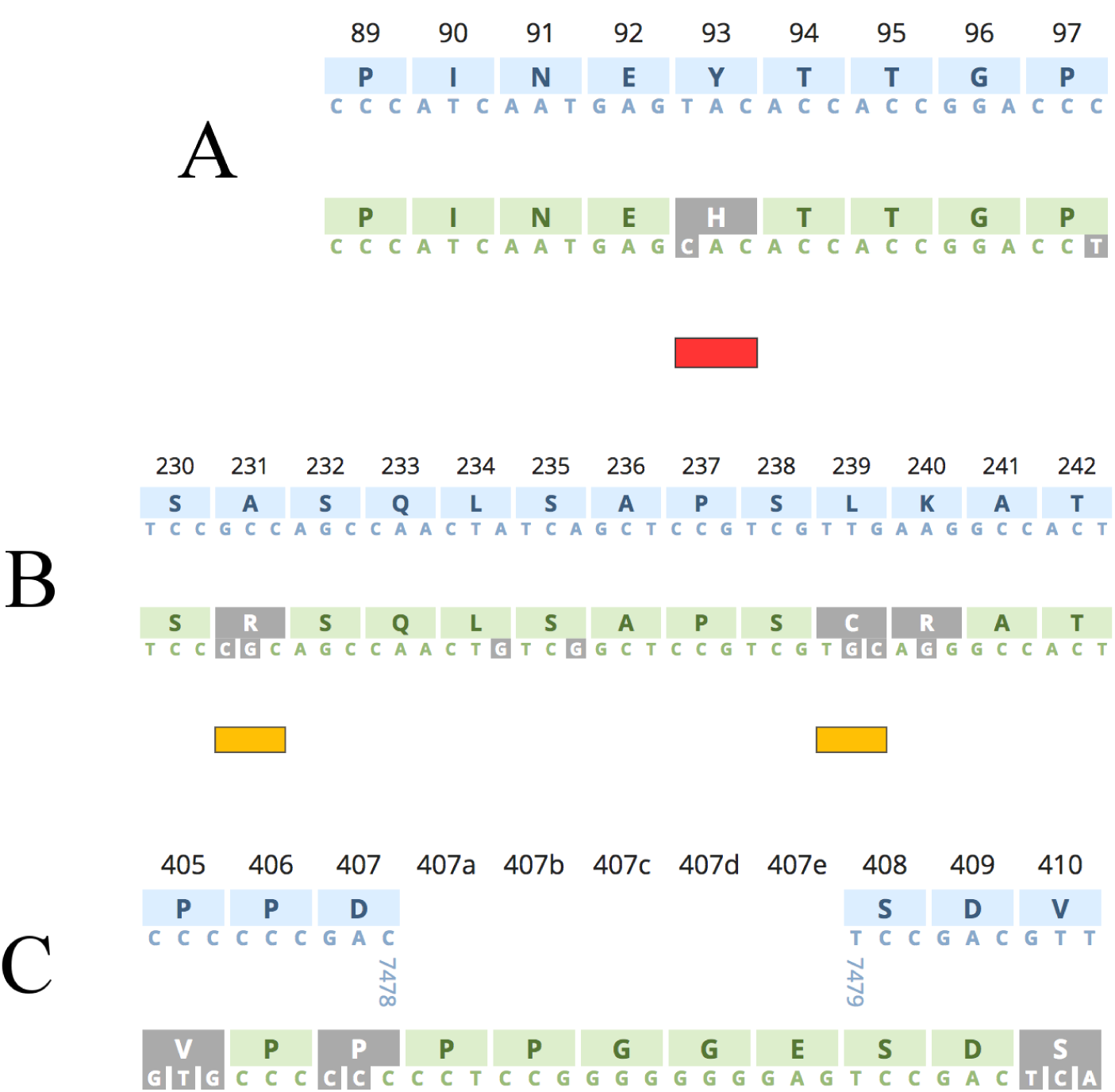
Genome visualisation of the NS5A viral protein within a subtype 3a query sequence via the HCV-GLUE web report. The submitted query sequence in green is contrasted with a user-selected reference sequence in blue. Differences between the query and reference are highlighted in inverted grey. In (A) the 93H RAS is highlighted by a red annotation bar and is shown to have arisen from a single nucleotide change relative to the subtype 3a reference (NC_009824) in the first nucleotide position within the codon. In (B) the query sequence contains amino-acid differences relative to the subtype 3a reference sequence at positions 231, 239 and 240. There are nucleotide differences within codons 234 and 235 but these are synonymous. The amino acid differences at 231 and 239 are atypical for subtype 3a as indicated by the yellow annotation bars, whereas the Arginine (R) at position 240 is typical for the subtype. In (C) the subtype 3a query sequence is shown alongside the HCV master reference (NC_004102); an insertion of 5 amino acids in the query sequence between codon locations 407 and 408 is shown; the consecutive nucleotide positions 7478 and 7479 in the master reference are also given to clarify the insertion

### Identification of resistance-associated substitutions

The high rates of successful cure achieved by DAA treatment have transformed the landscape for management and outlook for HCV-infected patients. However certain virus genome polymorphisms are associated with lower rates of sustained virological response (SVR). Therefore, there is considerable interest in identifying and monitoring polymorphisms that give resistance to DAAs from both the clinical and scientific communities.

At present, the HCV-GLUE dataset contains a catalogue of 162 resistance-associated substitutions (RASs), drawn from three recent survey articles [41, 42, 43]. These are represented using the GLUE *Variation* object type. Each RAS is defined by one or more virus genome locations and specific amino acid residues at these locations. Additionally, HCV-GLUE includes a schema extension which links each RAS *Variation* object with a characterisation of the evidence base for drug resistance, taken from the research literature and from current clinical guidance documents. This allows the web-based sequence interpretation system to present authoritative guidance and other relevant information about the RAS to the user (see figure 7). The aim will be to regularly update HCV-GLUE as guidelines become refined with options for DAA therapy and identification of any novel RAS.

**Figure 7.**
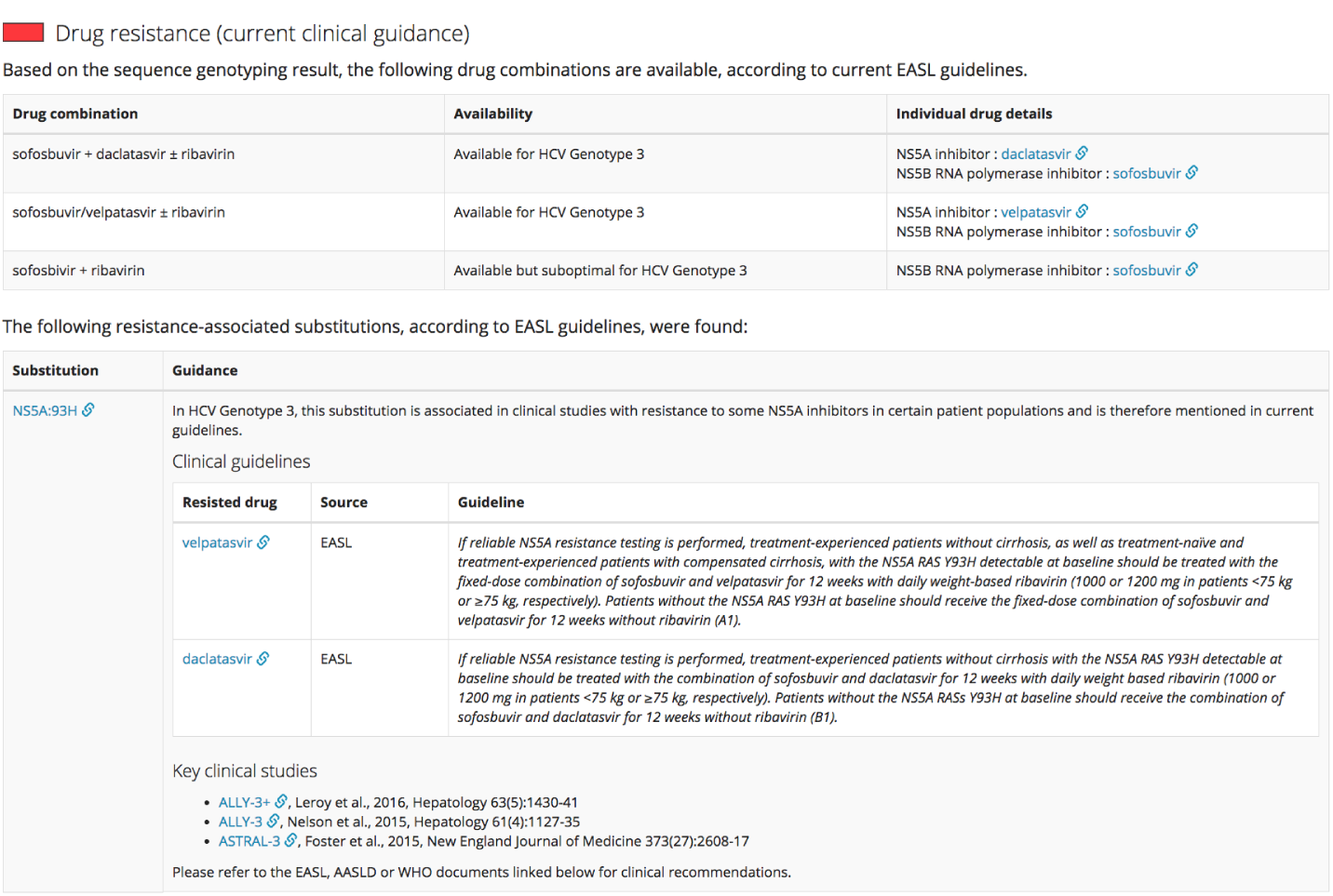
Web-based drug resistance report generated by HCV-GLUE. In this case the submitted sequence has been assigned to Genotype 3. The 93H substitution in the NS5A viral protein was detected; the report highlights that this is associated in the EASL guidance with resistance to both velpatasvir and daclatasvir.

### Comparison with other resources

Our case study resource HCV-GLUE can be compared with a number of existing resources. Sequence repository features similar to those of HCV-GLUE are found in the Los Alamos and ViPR HCV resources [7, 12]; sequences and metadata are extracted from GenBank and a clade assignment method is applied, subsets of sequences and alignments may be extracted based on various criteria. Both the ViPR HCV resource [7] and the REGA tools [1] provide a genotyping service for submitted sequences. HCV-GLUE clade assignment is comparable; on the one hand HCV-GLUE uses a slightly more sophisticated phylogenetic method but on the other hand it does not yet handle recombinant sequences. Drug resistance analysis for submitted sequences is provided by geno2pheno[hcv] [11] but has a slightly different emphasis from that of HCV-GLUE; geno2pheno[hcv] predicts the overall level of resistance of the virus to each drug based on the virus sequence and a set of rules. HCV-GLUE aims to highlight the evidence base relating to resistance-associated substitutions found within the virus sequence. One particular benefit of HCV-GLUE is that the entire GLUE project with all its data and analysis capabilities can be downloaded for use on a local computer.

## Further work

Interpretation of virus genome data often involves virtual translation of coding domains. In many viruses, such as HCV, this is straightforward to model. However, in some viruses, protein expression is complicated by a number of transcriptional and post-transcriptional phenomena, many documented in the ViralZone resource [44]. One example is transcriptional editing in the glycoprotein (GP) region of the Ebola virus genome. Another example is RNA splicing, for example in the tat protein in HIV-1. To model these phenomena in GLUE, the *Feature* and *FeatureLocation* object types could be extended in the core schema to allow transcriptional editing, RNA splicing and other phenomena to be specified in accordance with experimentally observed molecular processes.

GenBank sequence records have the capability of representing coding domains and other phenomena in the form of genome annotations. However these may be inconsistent across sequences even within the same virus species. Generating these annotations can be a significant bottleneck in the process of submitting sequences. GLUE is in a good position to help with the production of these annotations; if new sequences are stored in a GLUE alignment, GLUE may infer an annotation for the new sequence based on its homology with a reference sequence. GLUE is also a natural staging area for sequence metadata. We therefore think a new GLUE module type which generates GenBank submission files could improve the submission process.

Alignments are at the centre of any GLUE project, however these necessarily contain a level of uncertainty, with areas of poor quality. There are now a number of tools for assessing the quality of alignments and suggesting improvements, for example GUIDANCE2 [45] and trimAl [46]. These could be integrated into GLUE via new module types, allowing GLUE project developers to quickly establish a high-quality alignment before moving to downstream analysis.

The arrangement of sequences within a single alignment tree presumes a straightforward model of evolution in which different sections of the genome of a given virus have the same evolutionary history. This assumption is introduced for pragmatic reasons, as it is in many phylogenetic analyses of viruses. However, the welldocumented phenomena of of reassortment and recombination break this assumption.

So called “segmented” viruses (e.g. influenza, rotaviruses) possess a genome that is partitioned across physically distinct genomic segments. In these cases, multiple virus strains co-infecting the same cell can “reassort” and generate virus progeny that is distinct from the parental strains. In applying GLUE to such viruses we propose that each *Sequence* object should contain only the genetic material from one segment. A separate alignment tree should be introduced to capture the evolutionary hypothesis for each segment, containing only *AlignmentMember* objects for *Sequences* of that segment. *Sequence* objects representing different segments derived from the same biological sample can be linked via the use of schema extensions, which then facilitates analysis of reassortment.

Some viruses with non-segmented genomes (e.g. dengue, HIV-1) exhibit genetic recombination; within a host containing different strains, a new strain may appear containing genetic material from multiple “parent” virus strains in the same genomic fragment. In GLUE projects relating to viruses where recombination is occasional, this could be modelled within a single alignment tree. For a recombinant *Sequence*, multiple *AlignmentMember* objects would be created in order to assign different regions of the genome to different constrained *Alignments* reflecting the distinct origin of these regions. In GLUE projects where recombination is widespread, it may be more expedient to introduce separate alignment trees for those genes or genomic regions for which integrity tends to be preserved across recombination events.

## Discussion

The advent of powerful, affordable approaches for sequencing nucleic acids has exerted an enormous impact on virology and enabled new approaches for tackling viral disease. Accordingly, recent years have seen the emergence of a diverse range of databases and computational tools that are centred around virus genome data. These sequence data resources are being developed for a range of different reasons, in an environment characterised by rapid change and considerable uncertainty, and this creates numerous challenges and inefficiencies. In this paper, we have presented GLUE, a software system that has been designed specifically for the production of such resources.

The core schema of GLUE has been developed to capture the most important aspects of virus sequence data analysis. In particular, since comparative evolutionary analysis is typically central to sequence interpretation, GLUE places multiple sequence alignments at the centre of the data model. This means that common analysis functions such as variation scanning and clade assignment can be set up quickly for different virus projects.

To the best of our knowledge, GLUE is unique in terms of design and aims. The software underlying nextflu [9] has been generalised across multiple viruses [10] and is open source. It has a strong emphasis on real-time molecular epidemiology and graphical visualisation on the web, in contrast with the broader aims of GLUE. Some modules of the open source Chado project [21] overlap the core schema design of GLUE and the Chado schema could play a role in virus sequence data resources, since post-transcriptional phenomena such as splicing have been carefully modelled. However Chado only aims to solve the schema design problem while much additional functionality is required to build a resource such as HCV-GLUE.

By defining a restricted set of fundamental components, the core schema of GLUE introduces a degree of order and standardisation to the way in which sequence data resources are developed. Furthermore, each GLUE-based resource may define its own specific data extensions and analysis functionality, whilst also benefitting from the cross-cutting concerns addressed within the GLUE engine such as common bioinformatics functionality, database mechanisms, the interactive command line and web service integration. This high-level of flexibility should increase the versatility of GLUE as a system for constructing virus sequence data resources. These ideas have been validated in HCV-GLUE, a sequence data resource for hepatitis C virus with clinical and research applications.

## Conclusions

GLUE is a unified software environment supporting the rapid development of virus sequence data resources and promoting the efficient use and reuse of virus sequence expertise. We propose that GLUE can facilitate a step-change in the efficiency with which genomic data are used to advance research and public health.

## Availability and requirements

See the table below for details of the GLUE software availability and requirements.

|  |  |
| --- | --- |
| **Project name:** | GLUE (Genes Linked by Underlying Evolution) |
| **Project home page:** | http://tools.glue.cvr.ac.uk |
| **Operating system(s):** | Linux, Mac OSX, Windows |
| **Programming language:** | Java |
| **Other requirements:** | Java Platform Standard Edition 1.8.0 or later<br>MySQL Server Community Edition 5.7 or later<br>BLAST+ 2.2.31 or later<br>MAFFT 7.299b or later<br>RAxML 8.2.8 or later |
| **License:** | GNU Affero General Public License 3.0 |
| **Any restrictions to use by non-academics:** | None beyond license terms |

## List of abbreviations

API: Application programming interface
BLAST: Basic Local Alignment Search Tool
DAA: Direct-acting antiviral
DNA: Deoxyribonucleic acid
EASL: European Association for the Study of the Liver
EPA: Evolutionary Placement Algorithm
GLUE: Genes Linked by Underlying Evolution
GP: Glycoprotein
HCV: Hepatitis C virus
HIV: Human immunodeficiency virus
HTTP: HyperText Transfer Protocol
IEDB: Immune Epitope Database
JSON: JavaScript Object Notation
MAFFT: Multiple Alignment using Fast Fourier Transform
MLCA: Maximum Likelihood Clade Assignment
ORM: Object-relational mapping
RAS: Resistance-associated substitution
RAxML: Randomized Accelerated Maximum Likelihood
RNA: Ribonucleic acid
SQL: Structured query language
SVR: Sustained virological response
URL: Uniform resource locator
ViPR: Virus Pathogen Database and Analysis Resource
XML: Extensible Markup Language

## Declarations

Ethics approval and consent to participate

Not applicable

## Consent for publication

Not applicable

## Availability of data and material

The datasets generated and/or analysed during the current study are available as a MySQL database download from http://hcv.glue.cvr.ac.uk, or from the HCV-GLUE GitHub repository, access to which will be provided by the corresponding author on reasonable request.

## Authors’ contributions

JBS designed and wrote the software and its documentation and wrote the manuscript. ECT, JH and JM contributed to the specification of HCV-GLUE. ECT designed the antiviral resistance analysis. ECT and JM tested the HCV-GLUE web site. JH conceived the MLCA method. RJG conceived the GLUE concept, contributed to the design of the core schema, tested aspects of the GLUE implementation and supervised the project.

## Acknowledgements

The authors would like to express their gratitude to Ellie Barnes and other members of the STOP-HCV consortium for their valuable input into the development of HCV-GLUE, to Ru Xu of the Guangzhou Blood Center who contributed some metadata improvements for Genotype 6 sequences and to David Robertson and Massimo Palmarini who provided valuable feedback on the manuscript.

